# Topical sulforaphane exacerbates disease and promotes the accumulation of inflammatory CD8^+^ T cells in a dermatitis model

**DOI:** 10.64898/2026.09.11.751007

**Authors:** Anthony Tukanowicz-Hassett, Sagar P. Bapat, Yuichi Yokoyama, Taku Kambayashi

## Abstract

**Background:** Atopic dermatitis (AD) is a chronic inflammatory skin disease characterized by skin barrier dysfunction and immune activation. Antioxidants such as sulforaphane (SFN) have shown benefit in mouse models of AD and are being explored as topical therapies.

**Objectives:** We investigated the immune mechanisms through which topical SFN affects skin pathology in the MC903-induced AD-like dermatitis.

**Materials and Methods:** We used flow cytometry to characterize skin immune cell infiltration, activation, and cytokine production. Genetic and pharmacological loss-of-function studies were used to determine the contribution of specific immune cell populations and signaling pathways.

**Results:** Although topical application of SFN by itself caused no skin inflammation, it exacerbated MC903-induced dermatitis. Disease exacerbation was accompanied by robust T cell infiltration to skin, particularly by CD8^+^ T cells, and was significantly attenuated in TCRβ-deficient and β_2_m-deficient mice. In addition to a Type 2 inflammatory response, the skin CD8^+^ T cells exhibited a Type 1 inflammatory program characterized by IFNγ production that was largely absent among CD4^+^ T cells. Loss of IFNγ signaling attenuated SFN-mediated disease exacerbation. Blocking lymphocyte egress with FTY720 markedly reduced the SFN-induced increase in skin CD8^+^ T cells, suggesting that the increase was primarily dependent on recruitment from the periphery rather than local expansion.

**Conclusion:** Topical SFN worsens MC903-induced dermatitis through a mechanism involving the peripheral accumulation of IFNγ-producing CD8^+^ T cells in skin. These findings demonstrate that SFN can exert context-dependent pro-inflammatory effects and underscore the need to evaluate its effects carefully when considering topical medical or cosmetic applications.

## Introduction

Atopic dermatitis (AD) is one of the most common inflammatory skin diseases, and is characterized by skin barrier dysfunction, pruritus, and inflammation, which results in reduced quality of life for patients [1]. In response to barrier disruption and keratinocyte stress, epithelial cells release alarmins such as IL-25, IL-33, and thymic stromal lymphopoietin (TSLP), which promote type 2 inflammation by activating group 2 innate lymphoid cells (ILC2s) and recruiting Th2-polarized CD4^+^ T cells that produce IL-4, IL-5, and IL-13 [2–4].

While AD is classically described as a CD4^+^ T cell-driven disease, growing evidence suggests that CD8^+^ T cells also contribute to disease pathogenesis [5–7]. Notably, higher CD8^+^ T cell frequencies have been found to be associated with disease severity [8, 9]. Patients with severe AD exhibit expansion of skin-homing CD8^+^ T cell subsets, particularly Tc2 (CD8^+^ T cells that produce IL-4, IL-5, and IL-13) and Tc22 (CD8^+^ T cells that secrete IL-22) populations, analogous to CD4^+^ Th2 and Th22 subsets, respectively [10]. In lesional skin, CD8^+^ T cells have been identified as sources of inflammatory cytokines, including IL-13, IFNγ, IL-17, and IL-22, suggesting that they may participate in both type 2 and mixed inflammatory programs [11]. In an allergen-driven mouse model of AD, skin-infiltrating CD8^+^ T cells, rather than CD4^+^ T cells, have been shown to initiate lesion formation, implicating them as key effector cells in cutaneous inflammation [12]. Immunophenotypic heterogeneity beyond the type 2 pathway in AD has been associated with reduced responsiveness to dupilumab, a monoclonal antibody that blocks IL-4 and IL-13 signaling via inhibition of the IL4Rα [13, 14]. Despite emerging evidence that CD8^+^ T cells may contribute to disease severity, widely used mouse models of AD lack robust CD8^+^ T cell infiltration, mainly relying on Th2 CD4^+^ T cells or innate immune cells to drive disease [15–17]. While some models demonstrate increased CD8^+^ T cell infiltration, this response is either transient or its causal contribution to disease has not been established [12, 18, 19].

In this study, we investigated the effects of sulforaphane (SFN), a phytochemical derived from cruciferous vegetables, on the MC903-induced mouse model of AD. MC903 serves as a well-established system in which topical application of the vitamin D3 analogue calcipotriol (MC903) drives type 2 skin inflammation, recapitulating many of the clinical and immunological characteristics of acute human AD [16, 20]. SFN is a strong activator of the NRF2 antioxidant pathway and has been proposed as a potential therapeutic approach for AD based on its ability to enhance skin barrier gene expression, reduce oxidative stress, and dampen inflammatory signaling [21, 22]. However, here we report the unexpected finding that SFN exacerbates disease in the MC903 mouse model of AD. Topical application of SFN to MC903-treated ear skin of mice markedly increased ear swelling and immune cell infiltration relative to ear skin treated with MC903 alone, with notable enrichment of inflammatory cytokine-secreting CD8^+^ T cells. Loss-of-function studies suggested that these CD8^+^ T cells actively drive the worsened pathology. Together, these findings highlight the potential for SFN to elicit unexpected and context-dependent inflammatory responses in skin. Moreover, the addition of SFN to MC903 could be used as a platform to dissect the mechanisms underlying CD8^+^ T cell pathogenicity in AD.

## Materials and Methods

### Mice

C57BL/6 female mice were purchased from Charles River Laboratories. TCRβ KO, β_2_M KO, and MHCII KO mice were purchased from The Jackson Laboratories. GREAT, Smart13 (SM13), and Smart17A (SM17) reporter mice were kindly provided by Dr. Richard M. Locksley (University of California, San Francisco) and intercrossed to generate triple cytokine reporter mice on the C57BL/6 background. GREAT mice express enhanced yellow fluorescent protein (eYFP) from the endogenous Ifng locus [23], Smart13 mice express a nonsignaling human CD4 surface reporter from the endogenous Il13locus [24], and Smart17A mice express a truncated, nonsignaling human nerve growth factor receptor (hNGFR; CD271) surface reporter from the endogenous Il17a locus [25]. Each reporter is expressed through an internal ribosomal entry site introduced downstream of the corresponding cytokine coding sequence, preserving endogenous cytokine expression. These mice permit simultaneous flow cytometric assessment of IFNγ, IL-13, and IL-17A reporter expression in freshly isolated cells without ex vivo restimulation. B6 IFNγR1 KO mice were kindly provided by Dr. Igor Brodsky (University of Pennsylvania). Unless otherwise specified, all mice were 8 to 12 weeks old at the time of use, were housed in pathogen-free conditions, and were treated in strict compliance with the Institutional Animal Care and Use Committee regulations at the University of Pennsylvania.

### Treatment of Ear Skin with MC903 and SFN

Mice were treated on both ears with 10 μL of 200 μM MC903 (Tocris Bioscience) dissolved in EtOH or with EtOH vehicle for up to 12 consecutive days. Following MC903 treatment, mice were treated on both ears with 10 μL of SFN (20 mg/ml) or with EtOH vehicle three times a week during the 12 day treatment period. To evaluate skin inflammation, the thickness of both ears was measured using a dial thickness gauge and averaged. On the indicated days post-treatment, mice were euthanized, and tissues were collected for analysis.

### Cell Isolation from Ear Skin and Flow Cytometry

Mice were euthanized by CO₂ asphyxiation and ears were harvested. The dorsal and ventral sheets were separated and placed in digestion buffer with RPMI 1640 containing 0.25 mg/ml Liberase TL (Roche Diagnostics) and 0.5 mg/ml DNase I (Sigma-Aldrich) for 1.75 hours at 37°C, followed by mincing with scissors and mechanical dissociation by pipetting. Cell suspensions were passed through a 70 µm nylon mesh filter and resuspended in PBS. The cells were then stained with Live/Dead Near-IR to exclude dead cells, and Fc receptors were blocked with anti-CD16/32 (clone 2.4G2; BD Biosciences) for 15 minutes at 4°C. Cells were then stained with surface antibodies for 30 minutes at 4°C. Intracellular antigens were stained using the eBioscience™ Foxp3 Transcription Factor Staining Kit according to the manufacturer’s protocol. In some experiments, single-cell suspensions from ear skin were stimulated ex vivo in medium consisting of α-minimum essential medium (α-MEM) supplemented with 10% FBS, 1% penicillin-streptomycin-glutamine (PSG), 12.5 mM HEPES, and 22.7 µM β-mercaptoethanol (β-ME), containing 50 ng/mL PMA, 1 µg/mL ionomycin, and 5 µg/mL brefeldin A (BFA) for 4 hours at 37°C in 5% CO₂. Following stimulation, cells were processed for surface staining and intracellular cytokine staining using the BD Cytofix/Cytoperm Kit according to the manufacturer’s protocol. Flow cytometry was performed with a Symphony A3 Lite flow cytometer (BD Biosciences). Data were analyzed using FlowJo software (TreeStar).

### Skin Histology

Skin tissue was fixed overnight at 4°C in 4% paraformaldehyde (Thermo Fisher Scientific) prior to paraffin embedding. Tissues were sectioned at 5 µm, mounted onto glass slides, stained with hematoxylin and eosin (H&E) using standard protocols and imaged (Keyence BZ-X700). Tissue processing and H&E staining were performed by the Skin Biology and Disease Resource-Based Core (SBDRC) at the University of Pennsylvania. Inflammation was scored in a blinded manner by two independent observers based on immune cell infiltration (0, no infiltration; 1, mild; 2, moderate; 3, severe). Epidermal and dermal thickness were measured using ImageJ (NIH).

### FTY720 Treatment and Data Analysis

Mice were injected daily with FTY720 (Cayman Chemical) at a dose of 1 mg/kg or with vehicle (2.5% DMSO in PBS). To quantify the increase in skin CD8^+^ T cell numbers specifically induced by the addition of topical SFN, we subtracted the mean skin CD8^+^ T cell number in the MC903-alone group from the CD8^+^ T cell number in each SFN+MC903-treated mouse. This calculation was performed separately for the vehicle- and FTY720-treated groups.

### Statistical Analysis

Statistical tests for ear thickness and flow cytometry data were performed using GraphPad Prism software. Longitudinal ear-thickness measurements were analyzed by two-way repeated-measures ANOVA with treatment and time as factors. Welch’s two-tailed t test was used for comparisons between two independent groups in flow-cytometric and endpoint analyses. To compare ear thickness across mice of different genetic backgrounds, the mean ear thickness of MC903-alone-treated mice was subtracted from the ear thickness of each SFN+MC903-treated mouse of the same genetic background within each experiment.

## Results

### Topical SFN exacerbates ear swelling in the MC903-induced mouse model of dermatitis

SFN is a phytochemical derived from cruciferous vegetables that displays antioxidant activity [26, 27]. Given the role of oxidative stress in the pathogenesis of AD, we initially hypothesized that the addition of topical SFN would protect against disease in a mouse model of AD [28, 29]. To this end, we utilized the MC903-induced mouse model of AD, which involves once-a-day topical application of MC903 on ear skin over the course of 12 days. To examine the effect of SFN on MC903-induced AD, we additionally treated the ear skin 3 times/week (30 minutes after MC903 treatment) with either vehicle (EtOH) or SFN. Contrary to our original hypothesis, mice treated with SFN+MC903 exhibited significantly increased ear swelling compared to mice treated with vehicle+MC903 (**Figure 1A**), suggesting that SFN worsens MC903-induced AD. Consistent with this observation, histological analysis of ear skin tissue revealed that SFN augmented MC903-induced tissue pathology, which was quantified by skin immune cell infiltration (**Figure 1B-C**) and epidermal/dermal thickness (**Figure 1D**). To more closely examine the immune cell infiltration, flow cytometry of the digested ear tissue was performed. Compared with MC903 alone, SFN+MC903 significantly reduced eosinophil numbers, consistent with reduced type 2 inflammation, and produced a nonsignificant trend toward increased neutrophil numbers (**Figure 1E-F**). Neutrophils are most strongly associated with type 3 inflammation but can also participate in type 1 inflammatory responses. Importantly, mice treated with SFN alone did not exhibit any signs of inflammation and were similar to vehicle-only controls **(Figure S1)**. These data demonstrate that SFN exacerbates MC903-induced dermatitis while altering the composition of the inflammatory infiltrate.

**Figure 1.**
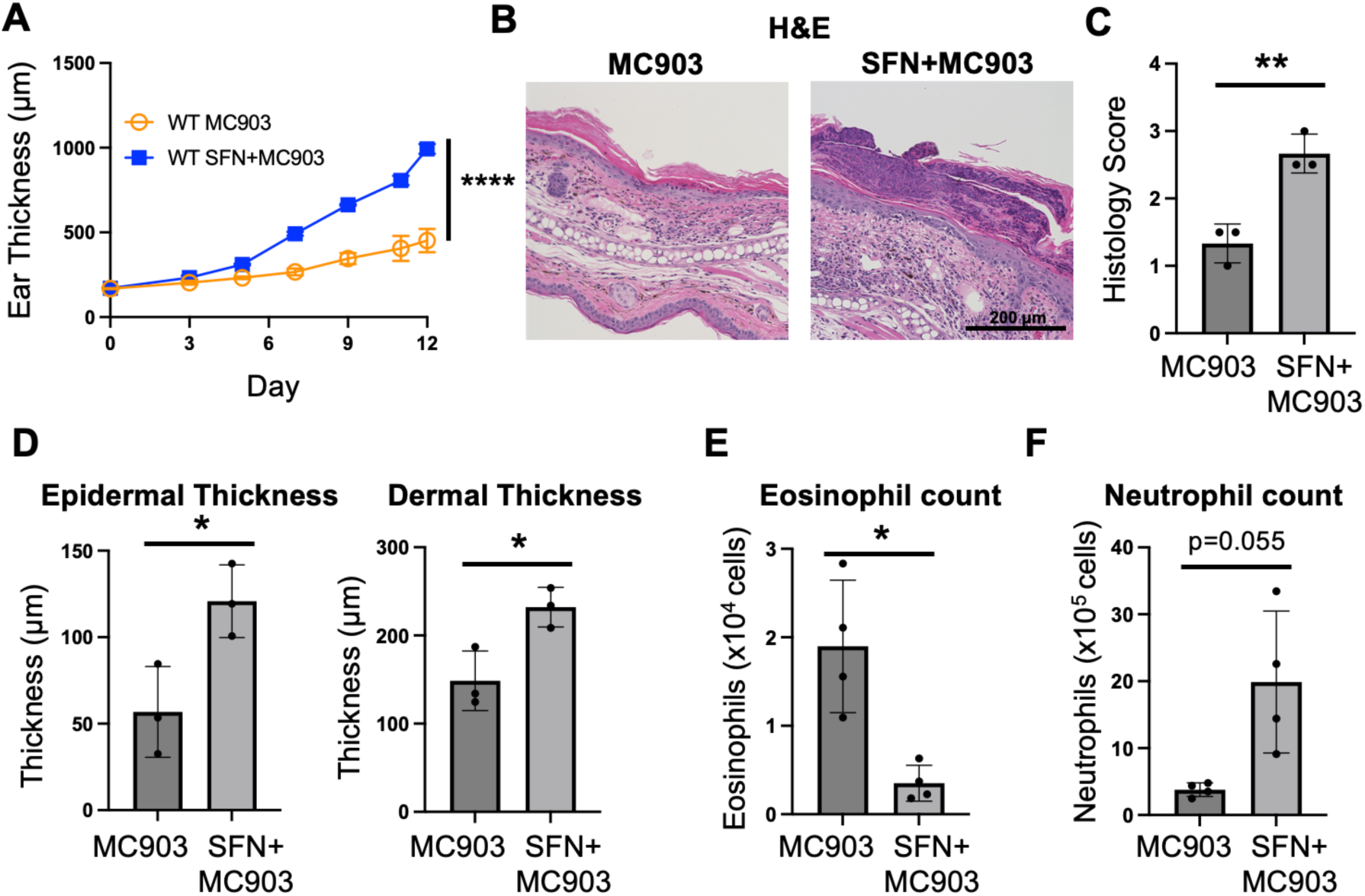
Topical SFN augments disease pathology in MC903-induced AD. Mice were treated daily with MC903 and three times weekly with vehicle or SFN for 12 days on ear skin. (**A**) Ear thickness was measured for 12 days. ****p<0.0001 by two-way ANOVA. (**B**) H&E staining of ear tissue, (**C**) compiled histopathological scores, and **(D)** epidermal and dermal thickness at Day 12 post MC903 and SFN+MC903 treatment are shown. **(E)** The number of eosinophils and (**F**) neutrophils in ear tissue of mice at Day 10 post MC903 and SFN+MC903 treatment was measured by flow cytometry. N = 3-4 mice/group/experiment. One representative of at least two independent experiments is shown. *p<0.05 and **p<0.01 by Welch’s t test.

### SFN-induced exacerbation of ear swelling is partially dependent on CD8^+^ T cells

Given that T cells are important drivers of AD pathology, we next examined whether SFN altered T cell populations in the ear skin of MC903-treated mice. Ear tissue from mice treated with SFN+MC903 contained significantly higher absolute numbers of both CD4^+^ and CD8^+^ T cells compared to mice treated with MC903 alone (**Figure 2A-B**). Although both T cell subsets increased, SFN treatment led to a disproportionate increase in CD8⁺ T cells, resulting in a relative enrichment of CD8⁺ T cells within the total TCRβ⁺ T cell compartment (**Figure 2C**).

**Figure 2.**
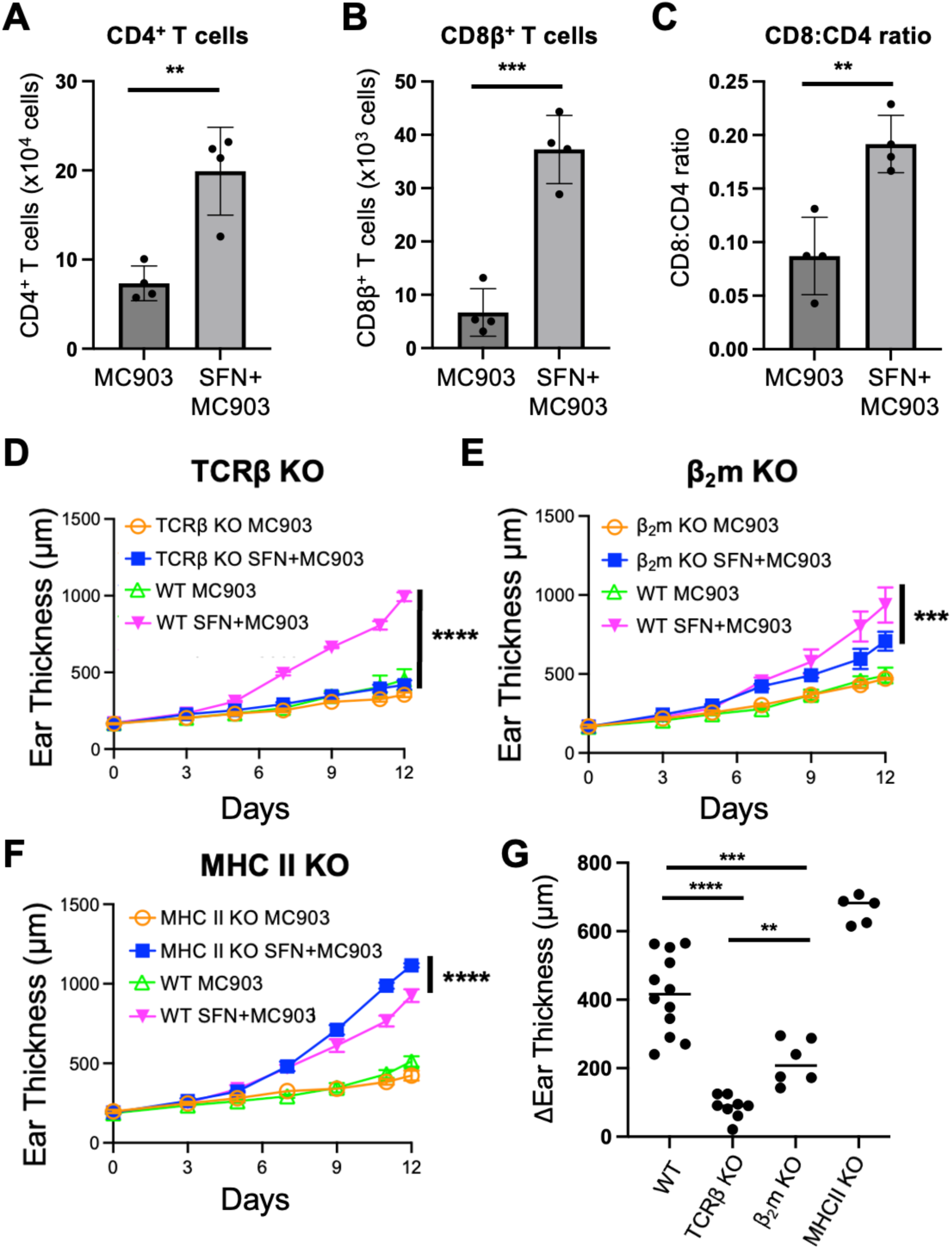
The exacerbation of MC903-induced inflammation by SFN is partially dependent on CD8^+^ T cells. Mice were treated daily with MC903 and three times weekly with vehicle or SFN on ear skin for 10 days. (**A)** CD4^+^ and **(B)** CD8^+^ T cell counts, and **(C)** the CD8:CD4 T cell ratio in ear tissue were determined by flow cytometric analysis. **p<.01 and ***p<0.001 by Welch’s t test. (**D**) The ears of TCRβ-deficient mice, **(E**) β2m-deficient mice (MHC I-deficient), (**F**) MHC-II-deficient mice, and respective WT control mice were treated with MC903 alone or with MC903+SFN for 12 days and ear thickness was measured. N = 3-4 mice/group/experiment. One representative of two independent experiments is shown. ***p<.001, ****p<0.0001 by two-way ANOVA. **(G)** Change in ear thickness of SFN+MC903-treated mice relative to the average ear thickness of MC903 alone within the same genetic background in each experiment. Each point represents an individual SFN+MC903-treated mouse pooled across experiments at Day 12 of treatment. **p<0.01, ***p<0.001, ****p<0.0001 via Welch’s t-Test.

To test whether the increase in skin T cells contributed to the SFN-mediated disease exacerbation, we subjected TCRβ-deficient mice (mice lacking TCRαβ T cells) to topical MC903 with or without SFN (**Figure 2D**). SFN+MC903-treated TCRβ-deficient mice exhibited a marked reduction in ear thickness compared to wild-type (WT) controls, indicating that TCRβ⁺ T cells are a major contributor to SFN-driven exacerbation of skin inflammation. To more specifically test the role of CD4^+^ and CD8^+^ T cell subsets in SFN-mediated disease exacerbation, we next assessed disease severity in β_2_m-deficient and MHC class II-deficient mice, which have profound deficiencies in conventional CD8^+^ and CD4^+^ T cells, respectively. Although not to the extent of TCRβ-deficient mice, β_2_m-deficient mice also displayed a significant reduction in the SFN-mediated increase in ear swelling, suggesting that CD8⁺ T cells played an important role in disease exacerbation (**Figure 2E**). Interestingly, however, MHC II-deficient mice treated with SFN+MC903 exhibited enhanced rather than reduced ear swelling compared to WT control mice (**Figure 2F**). Across multiple experiments, there was a statistically significant reduction in the SFN-mediated increase in ear thickness (difference in ear thickness of SFN+MC903 compared to MC903 alone) in TCRβ-deficient and β_2_m-deficient mice (**Figure 2G**).

### SFN leads to increased numbers of inflammatory cytokine-secreting CD8^+^ T cells

To understand which inflammatory programs were elicited in skin CD8⁺ T cells that were elicited by SFN+MC903, we next characterized the expression of lineage-determining transcription factors and cytokine production by skin CD8⁺ T cells (**Figure 3A-C**). For these analyses, we focused selectively on the skin CD8⁺ T cells that were increased in mice treated with SFN+MC903, since treatment with MC903 alone yielded too few CD8^+^ T cells for reliable analysis. In mice treated with SFN+MC903, CD8^+^ T cells expressing the transcription factors T-bet and GATA3, which drive Tc1 and Tc2 differentiation, respectively, were the most abundant CD8^+^ T cell subsets (**Figures 3A, S2A**). Tc17 cells, defined by RORγt expression, represented only a small fraction of all skin CD8⁺ T cells.

**Figure 3.**
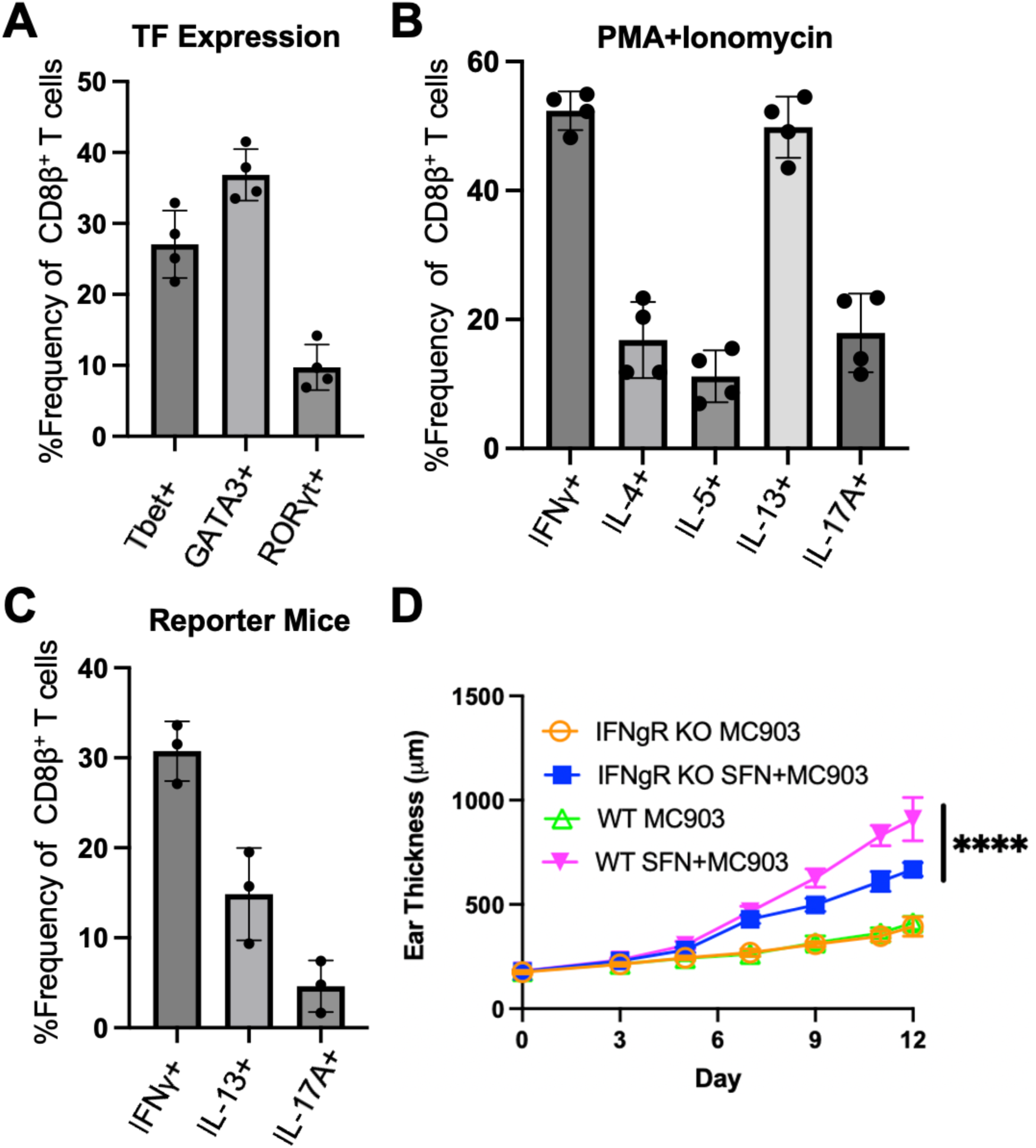
Topical SFN+MC903 treatment leads to increased inflammatory cytokine-producing CD8^+^ T cells in ear skin. Mice were treated daily with MC903 and three times weekly with vehicle or SFN on ear skin for 10 days. **(A**) The percentage of ear skin CD8^+^ T cells expressing Tbet, GATA3, and RORγt and (**B**) the indicated cytokines after PMA/ionomycin stimulation was determined by flow cytometry. (**C**) IFNγ/IL-13/IL-17A triple reporter mice were treated with MC903 and three times weekly with SFN on ear skin for 10 days. The frequency of freshly isolated CD8^+^ T cells expressing these reporters in ear skin was analyzed by flow cytometry. Data are shown as Mean ± S.D. for plots A-C. (**D**) IFNγR-deficient mice and WT control mice were treated daily with MC903 and three times weekly with vehicle or SFN on ear skin for 12 days. The mean ear thickness measurement ± S.D. is shown. N = 6-8 mice/group pooled from 2 independent experiments. ****p<0.0001 by two-way ANOVA.

We next investigated whether these CD8⁺ T cells were actively producing inflammatory cytokines. To assess the cytokine-producing potential of these CD8^+^ T cells, we restimulated skin T cells with PMA and ionomycin, followed by detection of intracellular cytokines by flow cytometry. Under these conditions, CD8^+^ T cells produced IFNγ (Tc1), IL-4/5/13 (Tc2), and IL-17A (Tc17), with IFNγ^+^ and IL-13^+^ CD8^+^ T cells being the most abundant (**Figures 3B, S2B-C**). To assess cytokine reporter expression without ex vivo restimulation, we analyzed CD8⁺ T cells freshly isolated from the skin of IFNγ/IL-13/IL-17A triple reporter mice. Consistent with their transcription factor profiles, these cells predominantly expressed the IFNγ and IL-13 reporters, whereas IL-17A reporter expression was limited (**Figures 3C, S2D**). Notably, IFNγ was minimally detected in CD4^+^ T cells (**Figure S3A-D**), suggesting that the Type 1 cytokine signature was largely restricted to the CD8^+^ T cell compartment.

Because IFNγ production was particularly prominent among skin CD8^+^ T cells, we next tested whether IFNγ signaling contributed to SFN-driven exacerbation. To this end, we treated the ears of IFNγR-deficient mice with MC903 with or without SFN and measured ear thickness over time (**Figure 3D**). When treated with SFN+MC903, IFNγR-deficient mice showed significantly reduced ear swelling compared to WT control mice, suggesting that IFNγ signaling plays an important role in SFN-mediated exacerbation of skin inflammation.

### The increase in skin CD8^+^ T cells induced by SFN+MC903 is primarily dependent on peripheral recruitment

We next sought to determine how CD8^+^ T cells were increased in the skin by SFN+MC903 treatment. Increased T cell numbers in inflamed skin could result from several mechanisms, including recruitment from the periphery or local expansion of tissue-resident cells. To test the contribution of these two mechanisms in the SFN-mediated increase in skin CD8^+^ T cells, we injected mice with either vehicle or FTY720, an S1P receptor modulator that limits lymphocyte egress from secondary lymphoid organs and thereby reduces circulating lymphocytes [30]. At the same time, we topically treated the ears of these mice with MC903 with or without SFN. As expected, FTY720 treatment induced marked peripheral lymphopenia, as demonstrated by reduced frequencies of TCRβ^+^ T cells and CD19^+^ B cells in blood (**Figures 4A-B**). Compared to vehicle control mice, FTY720-treated mice that were topically administered SFN+MC903 exhibited a significant reduction in skin CD8^+^ T cells (**Figure 4C**). To directly compare the increase in T cell numbers induced by the addition of SFN to MC903 in the presence or absence of FTY720, we calculated the difference in CD8^+^ T cell numbers between SFN+MC903 and MC903-treated mice within each group. This analysis revealed that FTY720 nearly completely reduced the SFN+MC903-induced increase in skin CD8^+^ T cells (**Figure 4D**). We next tested whether the residual CD8^+^ T cells in the skin after FTY720 treatment were increased due to proliferation by examining the expression of Ki-67, a nuclear protein that is expressed during the cell cycle. No relative differences in the frequency of Ki-67^+^ CD8^+^ T cells could be seen in groups that received SFN with or without MC903, indicating that the addition of SFN does not significantly promote the proliferation of tissue-resident CD8^+^ T cells (**Figures 4E-F**). Together, these findings indicate that the SFN-induced increase in skin CD8^+^ T cells is largely dependent on recruitment from the periphery rather than enhanced local proliferation.

**Figure 4.**
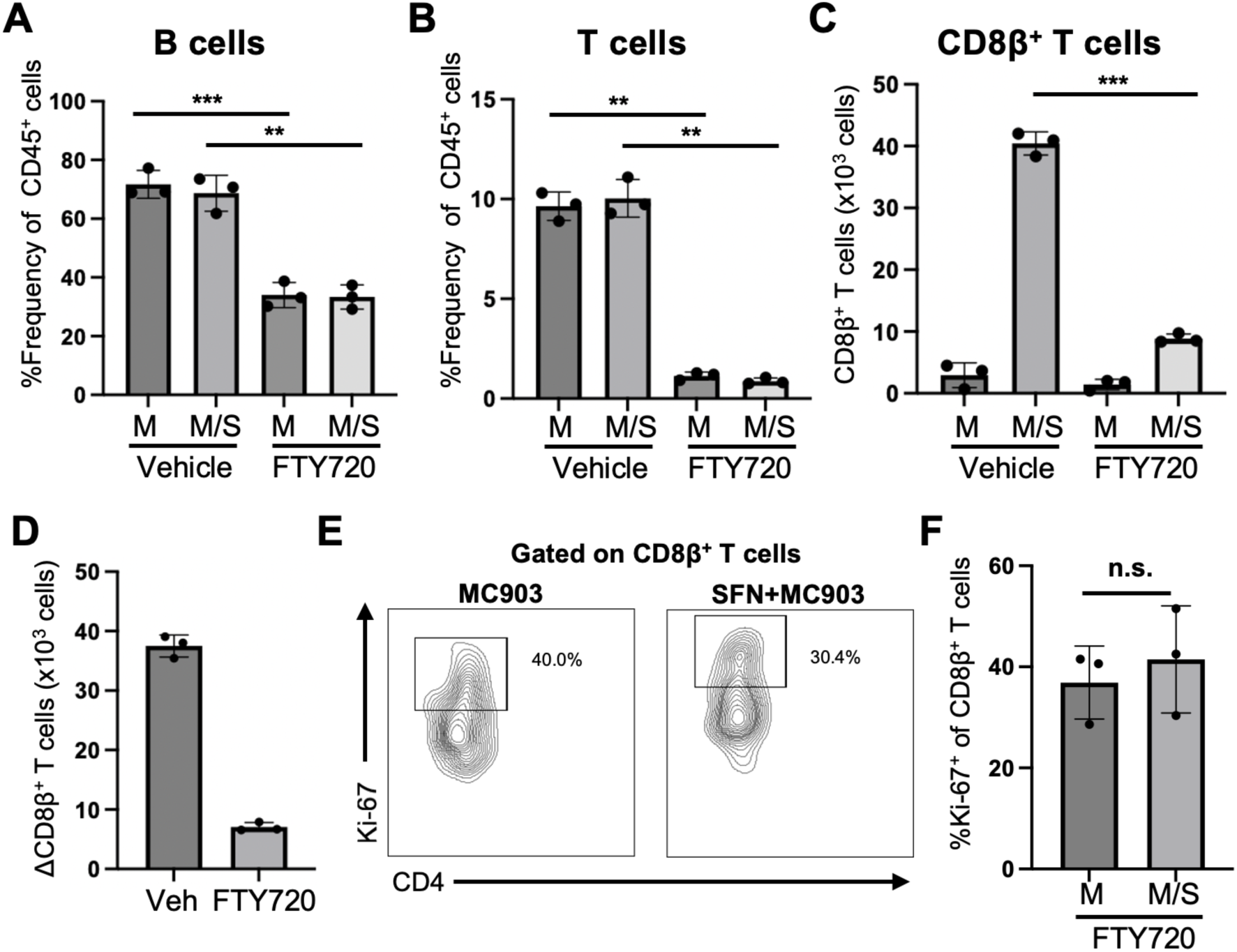
The increase in CD8^+^ T cells induced by SFN+MC903 is primarily driven by recruitment from the periphery. Mice were treated daily with vehicle control (saline) or FTY720. Two days after initial FTY720 treatment, mice were treated daily with MC903 and three times weekly with vehicle (M) or SFN (M/S) on ear skin for 10 days. (**A**)The frequency of B and **(B)** T cells amongst CD45^+^ cells in the peripheral blood was measured by flow cytometry. **(C)** The CD8^+^ T cell count in ear tissue was measured by flow cytometric analysis and the (**D**) difference in the number of CD8^+^ T cells in ears treated with SFN+MC903 compared to MC903-treated from either the vehicle control (Saline) or FTY720 treatment groups was calculated. (**E, F**) The frequency of Ki-67^+^ CD8^+^ T cells in the ears of FTY720-treated mice topically treated with MC903 or SFN+MC903 was measured by flow cytometry. ***p <0.001, **p<0.01 via Welch’s t-Test. Results are shown as Mean ± S.D. One representative of 2 independent experiments is shown.

## Discussion

In this study, we report an unexpected finding that topical SFN exacerbates MC903-induced skin inflammation. Notably, topical SFN in the presence of MC903 led to the disproportionate accumulation of skin CD8^+^ T cells that produced IFNγ. These CD8^+^ T cells contributed to disease exacerbation, since ear swelling was significantly reduced in T cell-deficient and β_2_m-deficient mice treated with MC903+SFN. Moreover, IFNγR-deficient mice exhibited partial attenuation of ear swelling, implicating IFNγ signaling as an important contributor to SFN-mediated pathology.

SFN activates NRF2 signaling through modification of KEAP1, reducing NRF2 degradation and promoting its nuclear accumulation and induction of cytoprotective antioxidant programs [21, 27]. SFN has demonstrated anti-inflammatory and cytoprotective effects in multiple preclinical settings, including pulmonary fibrosis, neuroinflammatory conditions, and UV-induced skin injury [31–33]. In contrast, we found that topical SFN exacerbated inflammation in the MC903 model, while SFN alone produced no detectable skin inflammation. These divergent effects highlight the context dependence of SFN biology and may reflect differences in route of administration, tissue environment, or the pre-existing inflammatory state. Thus, the effects of topical SFN cannot necessarily be predicted from its antioxidant activity in other settings and warrant evaluation in the specific inflammatory context in which it is used.

SFN+MC903 treatment led to increased accumulation of skin CD8^+^ T cells from the periphery. The reduction in SFN-mediated ear swelling in TCRβ- and β2m-deficient mice supports an important contribution of conventional CD8^+^ T cells to disease exacerbation. The reduction of ear swelling in β_2_m-deficient mice was less pronounced than that observed in TCRβ-deficient mice, potentially reflecting contributions from additional αβ T cell populations or broader immunologic differences between these strains. To examine the contribution of CD4+ T cells, we tested MHC II-deficient mice. In contrast to β_2_m deficiency, MHC II deficiency enhanced SFN+MC903-induced ear swelling. A similar pattern has been reported in a Der p sensitization model, in which CD8^+^ T cell ablation attenuated disease whereas CD4^+^ T cell deficiency worsened it [12]. One possibility is that loss of MHC II-restricted CD4^+^ regulatory T cells removes an immunosuppressive component of the response.

While AD has a strong Type 2 immunity component, growing evidence has underscored the importance of the Type 1 immune axis in driving chronic AD. During the acute phases of AD, Type 2 cytokines are most abundant in the skin, with relatively low levels of IFNγ. However, IFNγ production and Th1 involvement become increasingly more important as AD transitions into the chronic phases of disease, a shift that has also been shown in a house dust mite-induced allergen patch test [34, 35]. Clinically, chronic AD is characterized by scaly, lichenified plaques accompanied by epidermal hyperplasia [36]. This Type 1 skewing is also a hallmark of the intrinsic AD (iAD) endotype, which is characterized by elevated IFNγ^+^ T cell frequencies in both blood and lesional skin [36].

CD8^+^ T cells accumulating in SFN+MC903-treated skin displayed prominent Tc1- and Tc2-associated programs, including production of IFNγ and IL-13 In contrast, skin CD4+ T cells produced little IFNγ, indicating a CD8-biased Type 1 component to the inflammatory response. Genetic loss of IFNγR reduced disease severity in SFN+MC903-treated mice, demonstrating an important contribution of IFNγ signaling to the exacerbated pathology. IFNγ is known to reduce skin barrier integrity through multiple mechanisms, including altering the ceramide populations within the stratum corneum and downregulating claudin-1 to impair tight junction function [37, 38]. Thus, although the precise mechanism remains to be established, CD8^+^ T cell-derived IFNγ may contribute to barrier disruption and the exacerbated pathology observed in this model. Together, these findings suggest that SFN introduces a prominent Type 1 component into the otherwise Type 2-dominant MC903 response, potentially modeling aspects of the mixed inflammatory programs observed in chronic AD. The upstream mechanism by which SFN promotes CD8^+^ T cell accumulation remains unknown. Future studies will be needed to determine whether SFN alters keratinocyte-derived chemokines or other signals that promote recruitment of inflammatory CD8^+^ T cells to MC903-treated skin.

In conclusion, we have provided data showing that topical SFN worsens inflammation in the MC903 dermatitis model, with an important contribution from infiltrating CD8^+^ T cells. This model may provide a useful platform for investigating the underappreciated contribution of CD8^+^ T cells and Type 1 inflammatory programs to AD-like skin inflammation. More broadly, these findings highlight the context-dependent effects of SFN and the need to evaluate its immunologic consequences in specific inflammatory skin settings.

## Acknowledgments

We thank S. Prouty of the Cutaneous Phenomics and Transcriptomic Core at Penn SBDRC for assistance with histological processing and staining and N. Apenes of the Igor Brodsky lab for facilitating procurement of IFNγR1 KO mice.

## Abbreviations

AD: Atopic dermatitis
BFA: Brefeldin A
β2m: Beta-2 microglobulin
EtOH: Ethanol
H&E: Hematoxylin and eosin
IFNγ: Interferon gamma
IFNγR: Interferon gamma receptor
IL: Interleukin
IL4Rα: IL-4 receptor subunit alpha
ILC-2: Group 2 innate lymphoid cell
KEAP1: Kelch-like ECH-associated protein 1
KO: Knockout
MC903: Calcipotriol
MHC: Major histocompatibility complex
Nrf2: Nuclear factor erythroid 2–related factor 2
PBS: Phosphate-buffered saline
PMA: Phorbol 12-myristate 13-acetate
S1PR1: Sphingosine-1-phosphate receptor 1
SFN: Sulforaphane
TCRβ: T cell receptor beta
Tc1: Type 1 cytotoxic T cell
Tc2: Type 2 cytotoxic T cell
Tc17: Type 17 cytotoxic T cell
Tc22: Type 22 cytotoxic T cell
Th1: Type 1 helper T cell
Th17: Type 17 helper T cell
Th2: Type 2 helper T cell
WT: Wild-type

## Author Contributions

A.T-H.: study conceptualization, study design, experimental work, data interpretation, and manuscript writing. Y.Y.: study conceptualization, data interpretation and manuscript review. T.K.: study conceptualization, study design, data interpretation, funding acquisition, manuscript writing, and project supervision.

## Supporting Information

**Supplementary table 1.**

| **<u>Reagent</u>** | **<u>Source</u>** | **<u>Identifier</u>** |
| --- | --- | --- |
| Live/Dead Near-IR | ThermoFisher | Cat# L10119 |
| Anti-mouse CD45 Antibody<br>(Clone 30-F11), BUV395 | BD | Cat# 103147 |
| Anti-mouse TCRβ antibody<br>(clone H57-597), BUV737 | BD | Cat# 612821 |
| Anti-mouse TCRβ antibody<br>(clone H57-597), PE-Cy7 | BioLegend | Cat# 109222 |
| Anti-mouse CD4 antibody<br>(clone RM4-4), BUV805 | BD | Cat# 741913 |
| Anti-mouse GATA3 antibody<br>(clone L50-823), PE | BD | Cat# 560074 |
| Anti-mouse RORγt antibody<br>(clone Q31-378), PE-CF594 | BD | Cat# 562684 |
| Anti-mouse T-bet antibody<br>(clone 4B10), PE-Cy7 | BioLegend | Cat# 644824 |
| Anti-mouse Foxp3 antibody<br>(clone FJK-16s), APC | eBioscience | Cat# 17-5773-82 |
| Anti-mouse CD8β antibody<br>(clone YTS156.7.7), Alexa<br>Fluor 700 | BioLegend | Cat# 126613 |
| Anti-mouse CD11c antibody<br>(clone N419), BV711 | BioLegend | Cat# 117349 |
| Anti-mouse CD11b antibody<br>(clone M1/70), FITC | BioLegend | Cat# 101206 |
| Anti-mouse CD11b antibody<br>(clone M1/70 ), PE-Cy7 | Biolegend | Cat# 101216 |
| Anti-mouse Ly6G antibody<br>(clone 1A8), PerCP-Cy5.5 | BioLegend | Cat# 127616 |
| Anti-mouse SiglecF antibody<br>(clone E50-2440), PE-CF594 | BD | Cat# 562757 |
| Anti-mouse SiglecF antibody<br>(clone E50-2440) | BD | Cat# 562680 |
| Anti-mouse CD90.2 antibody<br>(clone 53-2.1) | BioLegend | Cat# 40304 |
| Anti-mouse IFN-γ antibody<br>(clone XMG1.2) | BioLegend | Cat# 505830 |
| Anti-mouse IL-13 antibody<br>(clone eBio13A),<br>PerCPeFluor710 | eBioscience | Cat# 46-7133-82 |
| Anti-mouse IL-4 antibody<br>(clone 11B11), PE | ThermoFisher | Cat# 12-7041-82 |
| Anti-mouse IL-5 antibody<br>(clone TRFK5), APC | BioLegend | Cat# 504306 |
| Anti-mouse IL-17A antibody<br>(clone TC11-18H10.1), PE-Cy7 | BioLegend | Cat# 506922 |
| Anti-human NGFR antibody<br>(clone ME20.4) PerCP-Cy5.5 | BioLegend | Cat# 345112 |
| Anti-human CD4 antibody<br>(clone A161A1), APC | BioLegend | Cat# 357408 |
| Anti-mouse CD19 antibody<br>(clone 6D5), PE-Cy7 | BioLegend | Cat# 115520 |
| Anti-mouse Ki-67 antibody<br>(clone SolA15),<br>PerCPeFluor710 | eBioscience | Cat# 46-5698-80 |
| Anti-mouse Ki-67 antibody<br>(clone SolA15) FITC | eBioscience | Cat# 11-5698-80 |

**Supporting Information Figure 1.**
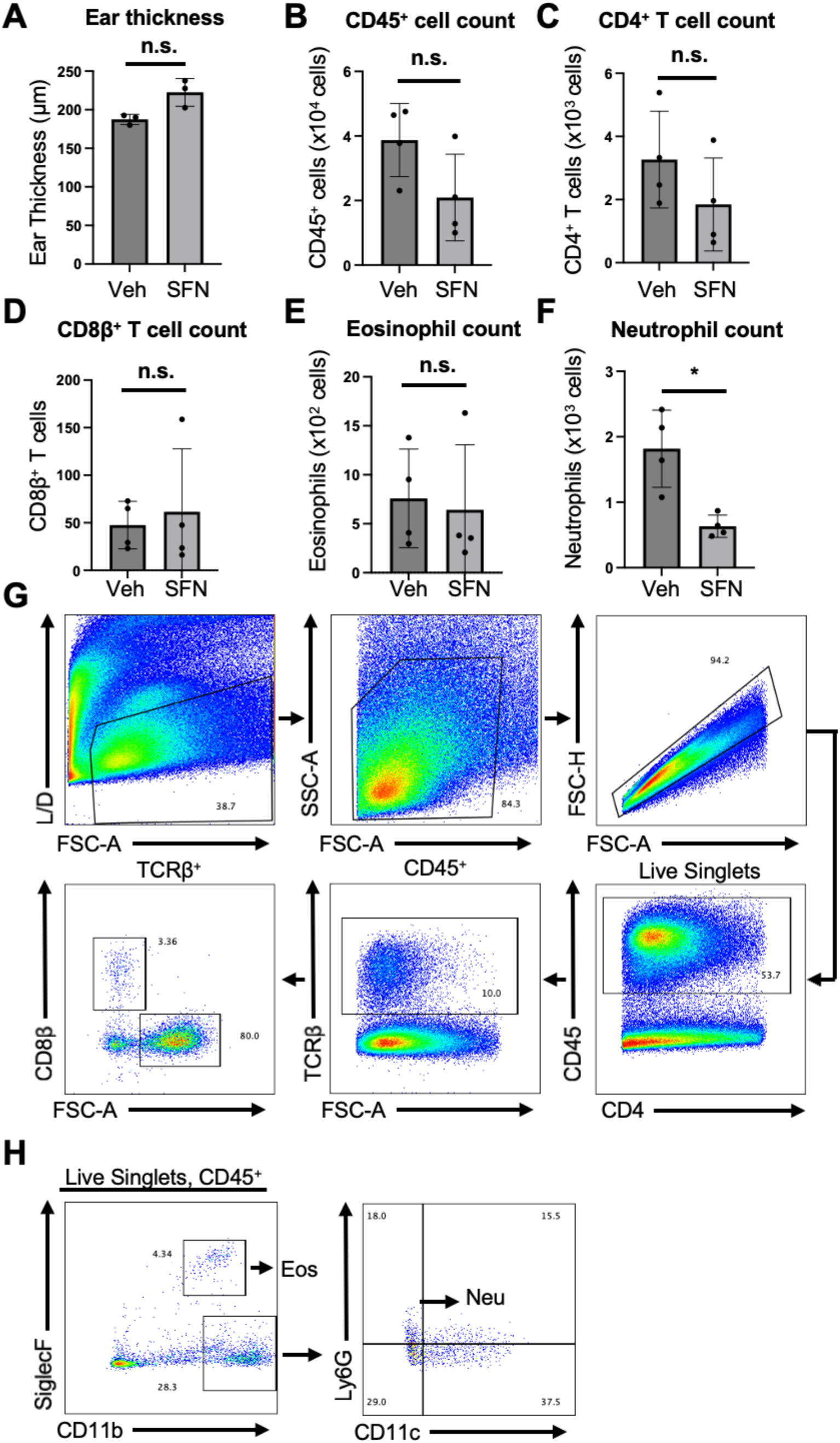
**(A)** Ear skin of B6 mice was treated daily with vehicle (EtOH) or with SFN for 12 days. Ear thickness on Day 12 is plotted as mean ± S.D. (n = 4). ns = not significant by Welch’s t-test. **(B)** Ear skin of B6 mice was treated daily with vehicle (EtOH) or with SFN for 6 days and the ear skin cells were analyzed by flow cytometric analysis for absolute numbers of CD45^+^ cells, **(C)** CD4^+^ T cells, **(D)** CD8^+^ T cells, **(E)** eosinophils, and **(F)** neutrophils. n.s. = not significant, *p<0.05 by Welch’s t-test. **(G)** Gating strategy used to identify lymphocyte and **(H)** myeloid populations by flow cytometry.

**Supporting Information Figure 2.**
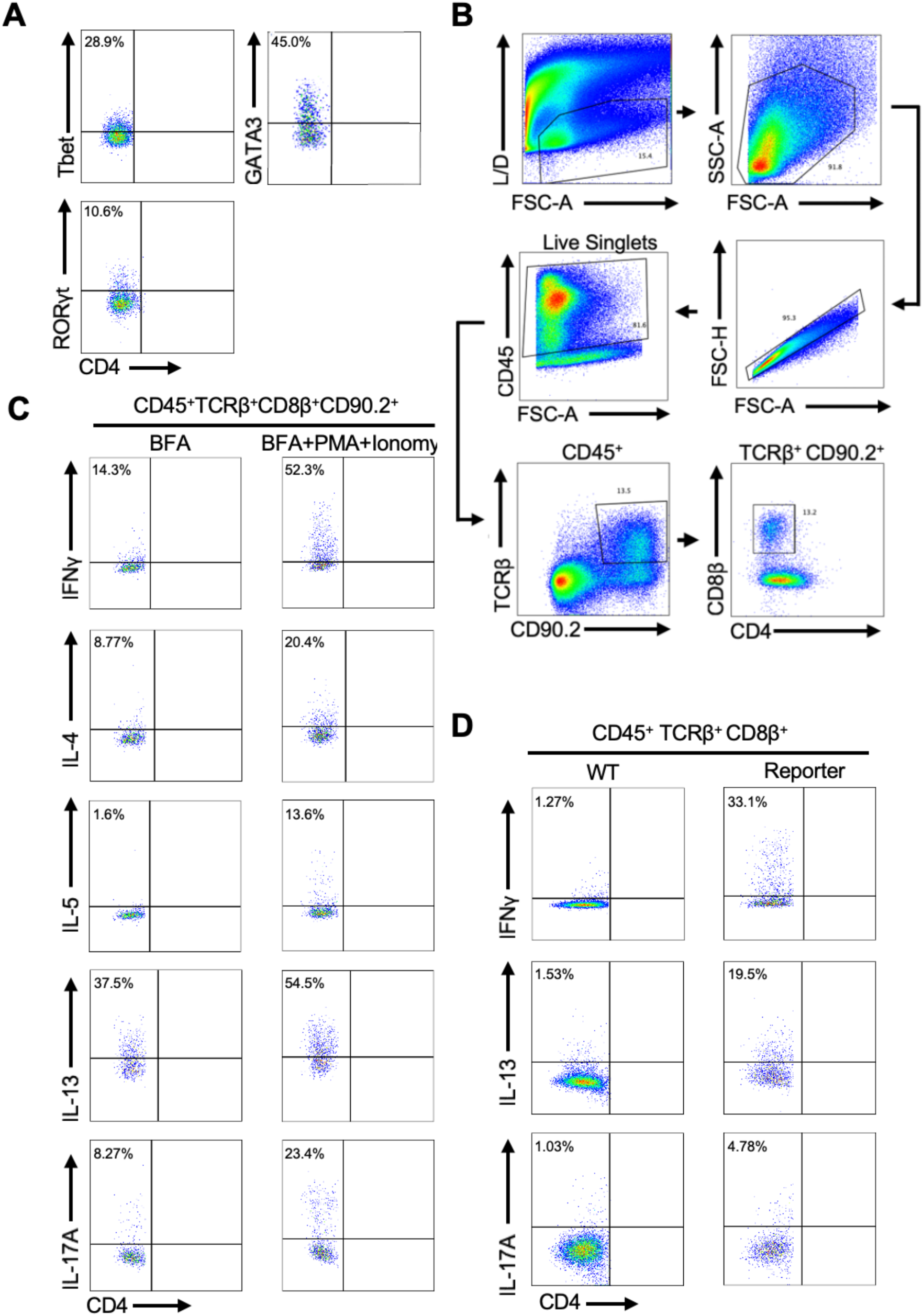
**(A)** Representative flow plots showing Tbet, GATA3, and RORγt expression among CD8β⁺ T cells from SFN+MC903-treated B6 mice. **(B)** Gating strategy for cells stimulated ex vivo with PMA/ionomycin. **(C)** Representative flow cytometry plots showing IL-4, IL-5, IL-13, IL-17A, and IFNγ expression by CD8β⁺ T cells isolated from the ears of B6 mice topically treated with SFN+MC903. Cells were analyzed following ex vivo with (right plots) or without (left plots) stimulation with PMA and ionomycin. (**D**) Representative flow cytometry plots showing IL-13, IL-17A, and IFNγ reporter expression among CD8β⁺ T cells from SFN+MC903-treated triple cytokine reporter mice (right plots). WT B6 mice (left plots) were included as negative gating controls. All plots are representative of CD8β⁺ T cells isolated from ear tissue on Day 10 of treatment.

**Supporting Information Figure 3.**
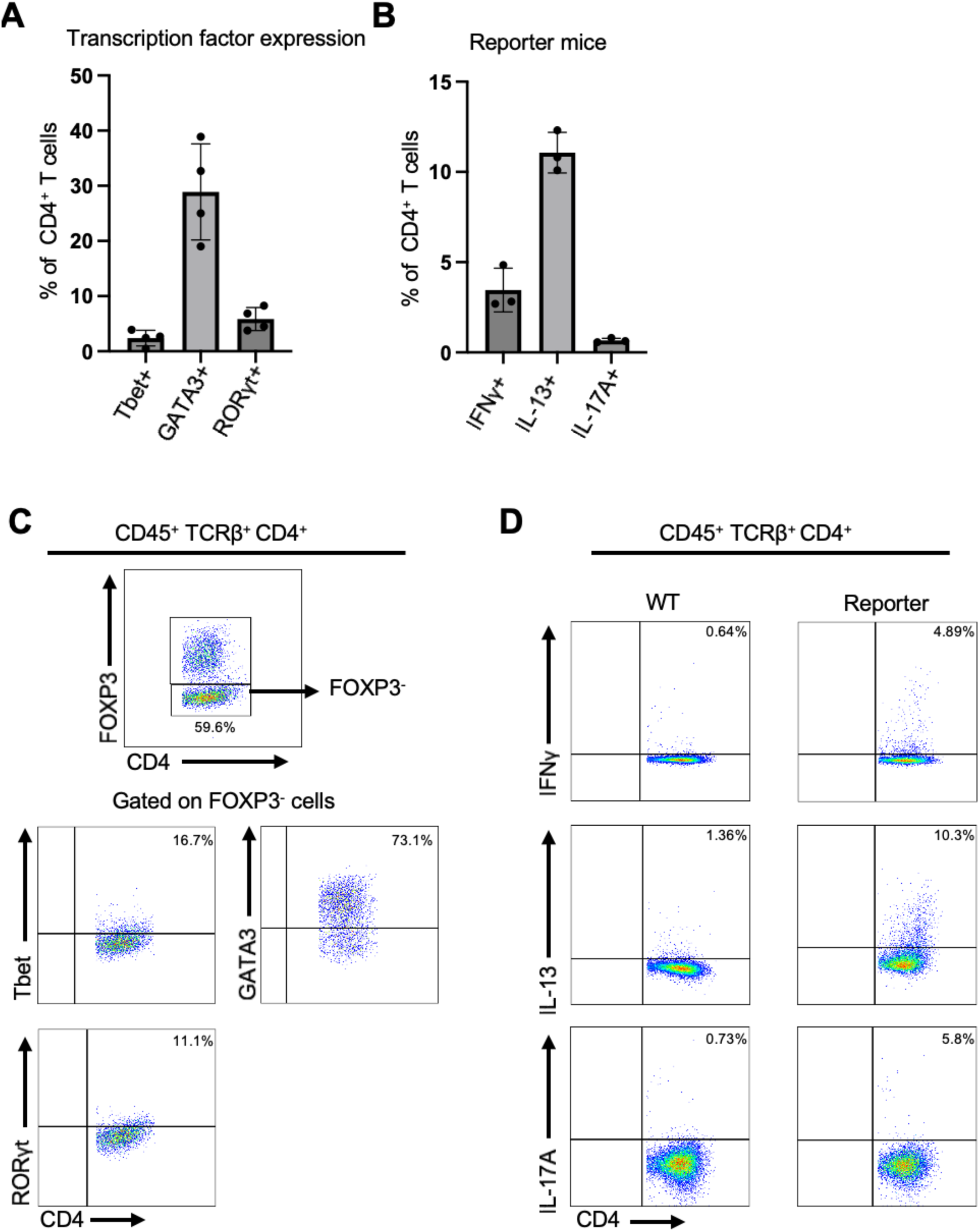
Ear skin of mice from WT and triple cytokine reporter mice were treated daily with SFN+MC903 and harvested on Day 10 for analysis of ear skin cells by flow cytometry. **(A**) The frequency of CD4^+^ T cells expressing GATA3, RORγt, or Tbet and (**B**) the frequency of CD4^+^ T cells expressing IL-13 (hCD4), IL-17A (NGF), or IFNγ (YFP) are shown as mean ± S.D. is shown in all plots. (**C**) Gating strategy and representative flow cytometry plots for Tbet, GATA3, and RORγt factor expression among CD4^+^ T cells. (**D**) Representative flow cytometry plots showing IL-13, IL-17A, and IFNγ reporter expression among CD4⁺ T cells from SFN+MC903-treated triple cytokine reporter mice (right plots). WT B6 mice (left plots) were included as negative gating controls.

## Notes

**Funding:** This research was supported by grants from the National Institutes of Health (R01AR082256-03).

**Conflict of interest:** The authors have no conflict of interest to declare.

### Competing Interest Statement

The authors have declared no competing interest.

